# Actively cycling cells in uninjured connective tissue are not a prerequisite for appendage regeneration

**DOI:** 10.64898/2026.07.25.740716

**Authors:** Emilio A. Oviedo-Rivadeneira, Ashley W. Seifert

**Affiliations:** Department of Biology, University of Kentucky, Lexington, KY 40506, USA; Department of Veterinary Anatomy and Physiology, University of Nairobi, Nairobi, Kenya

**Keywords:** Cell proliferation, regeneration, cell cycle

## Abstract

Multiple hypotheses have been formulated to explain differences in tissue repair ability across vertebrates. One hypothesis posits that the accessibility of actively cycling stromal cells within uninjured tissue confers access to a proliferative population in response to tissue damage. This hypothesis further suggests that animals with an indeterminate growth mode possess an actively cycling cell population necessary for growth that can be readily accessed for tissue regeneration. Moreover, the absence of an actively cycling population in connective tissue provides a mechanism that restricts regeneration in animals with determinate growth whose cells are refractory to cell cycle progression and proliferation to produce new tissue for morphogenesis. Here, we explore this paradigm using an EdU-BrdU pulse chase strategy in four different vertebrate species: two with determinate (*Acomys dimidiatus* and *Mus musculus*) and two with indeterminate modes of growth (*Danio rerio* and *Ambystoma mexicanum*). We find that although indeterminate growers do possess a small population of actively cycling cells, this population does not contribute to regeneration. Moreover, we found that while *Acomys* does not possess a population of actively cycling stromal cells, cells re-enter the cell cycle *de novo* in these animals to contribute to regeneration. Furthermore, testing this hypothesis allowed us to ask whether tissue injury could stimulate cell cycle re-entry – a so-called *primed* state - in cells at distance from the injury site in these four species and we did not find evidence of such priming in stromal or epidermal tissue.

**Highlights:**

- Cell cycle re-entry is a common response to injury in regenerative and non-regenerative vertebrates that is independent of actively cycling stromal cells in uninjured connective tissue
- Actively cycling cells do not contribute to regenerative healing in spiny mice, axolotls or zebrafish.
- Our data do not support systemic cell cycle activation in response to injury.

## INTRODUCTION

Nearly all species capable of complex tissue regeneration use epimorphic regeneration to rebuild missing tissue. Epimorphic regeneration is characterized by the proliferation and accumulation of a heterogeneous cell mass (aka a blastema) rather than transdifferentiation of resident cells to replace missing tissue (Morgan, 1901). Thus, during epimorphic regeneration, cell cycle entry must be initiated with cell cycle progression and division maintained to form enough new tissue to undergo morphogenesis.

To partially explain why some animals regenerate and others do not, scientists have formulated hypotheses regarding the ability of animal cells to maintain a proliferative response throughout regeneration. In contrast to highly regenerative animals, poorly regenerating species (e.g., holometabolous insects, most mammals, etc.) cannot maintain a proliferative response after injury. Some mammalian stromal cells and myocytes can re-enter the cell cycle in response to injury, but activation of certain tumor suppressors quickly arrests cell cycle progression (Gawriluk et al., 2016; Pajcini et al., 2010). The inability of mammals to maintain cell cycle progression has been hypothesized to underlie poor regeneration of complex tissues.

Additional hypotheses have been formulated to explain a breakdown in cell cycle progression experienced after tissue injury in mammals. Although only supported by correlative data, one hypothesis relates regenerative ability to growth mode (Pajcini et al., 2010; Pomerantz & Blau, 2013; Seifert, et al., 2012; Simkin & Seifert, 2018). Animals can grow determinately or indeterminately (Stearns, 1998) and determinate growers are so named because they stop growing after reaching a defined size range in a species-specific manner (Vinicius & Mumby, 2013). Vertebrates with determinate growth (e.g., mammals, birds, etc.) deploy relatively more cell cycle control mechanisms in comparison to vertebrates with indeterminate growth (salamanders, fish, etc.,) (Hesse et al., 2015; Pajcini et al., 2010; Pomerantz & Blau, 2013; Simkin & Seifert, 2018). For example, cell culture experiments using newt myotubes demonstrate that these cells easily re-enter the cell cycle after serum stimulation, whereas mouse myotubes formed from C2C12 cells are refractory to cell cycle re-entry (Tanaka et al., 1997; Tanaka & Brockes, 1998). Although *Mus* myotubes can be forced to re-enter the cell cycle, this requires deletion or inactivation of the tumor suppressors retinoblastoma protein (pRB) and p16^ink4a^ (Gu et al., 1993; Pajcini et al., 2010; Schneider et al., 1994). Similarly, the introduction of previously absent human *ARF* into zebrafish inhibits regeneration (Hesse et al., 2015). Differential regulation of cell cycle progression following injury in indeterminate growers and the fact that these animals require a population of cycling cells to sustain growth has led to the idea that the presence of this population could be one factor contributing to a highly regenerative phenotype (Pomerantz & Blau, 2013; Simkin & Seifert, 2018).

Here we explore several of these interconnected hypotheses, specifically the source of proliferating cells that participate in complex tissue regeneration and how progenitor cells are activated. First, we tested whether three highly regenerative vertebrate species possess actively cycling stromal cells that can support growth and tissue homeostasis. Next, we tested whether these proliferating stromal cells detected prior to injury are recruited for blastema formation and regeneration. Lastly, we tested if injury stimulates cell cycle re-entry in cells close to the injury and at distance (i.e., in contralateral tissue on the other side of the animal). We used a pulse-chase strategy to test these hypotheses in two mammals with determinate growth: one with enhanced regeneration abilities (*Acomys dimidiatus*) (Gawriluk et al., 2016; Riddell et al., 2025; Seifert, et al., 2012; Tomasso et al., 2023), and in one with generally poor regenerative ability (*Mus musculus*). Additionally, we investigated these hypotheses in two highly regenerative species with indeterminate growth modes (*Ambystoma mexicanum* and *Danio rerio*). Using this approach, we found that both species with indeterminate growth modes possessed a population of actively cycling stromal cells during tissue homeostasis. Surprisingly, this population did not contribute to regeneration during the early wound healing phase, at least in *Danio*. In contrast, we did not observe cycling stromal cells in the two species with determinate growth. However, stromal cells re-entered the cell cycle in both mammalian species in response to injury. Finally, we failed to find evidence for long-range cell cycle activation among stromal cells in any of the species studied.

## METHODS

### Animal care

Sexually mature, spiny mice (*Acomys dimidiatus* - aged six months to one year old), *Mus musculus*, (∼three months old) and *Ambystoma mexicanum* (subadults, 20-30 cm, and ∼65 grams) were obtained from our in-house breeding colonies at the University of Kentucky. Sexually mature *Danio rerio* were purchased and bred at a zebrafish facility at the University of Kentucky. Animals were maintained on a 12:12 h L:D light cycle. Rodent species were fed 14% mouse chow (Teklad Global 2014, Harlan Laboratories) and *Acomys* diets were supplemented with black sunflower seeds. Axolotls were fed salmon pellets and zebrafish with brine shrimp. All animal experiments were approved by the University of Kentucky Institutional Animal Care and Use Committee (IACUC) under protocol 2013–3254.

### Regeneration assays

Mammals were anesthetized with 3% vaporized isoflurane (v/v) (Henry Schein Animal Health, Dublin, OH) at 1 psi oxygen flow rate. A 4mm biopsy punch (Sklar Instruments, West Chester, PA) was used to create a through-and-through hole in the right and left ear pinna. This biopsy size has previously been shown to serve as a tissue regeneration assay in mammals (Gawriluk et al., 2016). Ear tissue was collected at specified time points using an 8mm biopsy punch circumscribing the original injury. Zebrafish were anesthetized in Tricaine-S (4mg/100ml) diluted in water (a.k.a., MS-222) and a blade was used to amputate caudal and pelvic fins through the fin rays. Animals were sacrificed after tissue collection. Axolotls were anesthetized with benzocaine 0.03% diluted in water and scissors were used to amputate and collect anterior limbs. The published protocol can be found here: dx.doi.org/10.17504/protocols.io.bp2l6dx65vqe/v1.

### Pulse-chase experiments

Ethynyl deoxyuridine (EdU) (10 ug/g) and Bromodeoxyuridine (BrdU) (1 ug/g) were diluted in normal saline solution for pulse-chase experiments (the published protocol can be found here: dx.doi.org/10.17504/protocols.io.261gerdmol47/v1). Both solutions were injected intraperitoneally every three hours over a 9-hr period in *Acomys*, *Mus* and *Ambystoma*. For *Danio*, solutions were diluted in the water tank for a 9-hr period. Animals were exposed to EdU before injury and to BrdU one or three days after injury. The left ear pinna in *Acomys* and *Mus,* the left limb in *Ambystoma*, and the caudal fin in *Danio* were collected after the EdU pulse to assess baseline cell numbers in S-phase, and the right pinna, right limb and pelvic fin respectively were collected after the BrdU pulse to track single (EdU+ or BrdU+) and double positive cells.

### Histology, EdU detection and Immunohistochemistry

Harvested tissue was fixed overnight in 10% neutral buffered formalin (NBF) (Thermo Fisher Scientific Inc. Epredia 5701) at 4°C on a shaker. After 16-20 hrs, the tissue was washed 3x with PBS, 3 minutes per wash, followed by 3x 70% ethanol washes for dehydration and storage at 4°C. Stored tissue was processed for paraffin (Leica Biosystems, Buffalo Grove, IL, Formula “R” REF: 3801450) embedding using a Histo5 tissue processor machine (Milestone). Samples were sectioned at 5μm. EdU was detected using click chemistry as follows: Tissue sections were de-paraffinized and rehydrated. Antigen retrieval was performed if co-stained with antibodies by exposure to high heat antigen retrieval in a steamer in antigen unmasking TRIS or citric acid-based solutions (Vector Laboratories, Inc.). Tissues were washed two times for 3 minutes in Tris-buffered saline (TBS) and incubated in the prepared reaction cocktail (see table below) for 30 minutes at RT protected from light. Tissues were washed two times in TBS for 3 minutes and if co-staining with antibodies was performed, sections were processed following the fluorescent IHC protocol for BrdU and cell type specific antibodies. Briefly, after a 3-minute wash, tissues were blocked in blocking serum (see table below) for 30 minutes at RT. Tissues were then incubated with primary antibody diluted in blocking serum at 4 °C overnight. After a 3-minute wash, tissues were incubated in secondary antibody diluted in blocking serum for 30 min at RT. Tissue samples were incubated with Hoechst 33342 for 5 minutes at RT and then dried and mounted in Prolong. For complete protocol details regarding paraffin sections (dx.doi.org/10.17504/protocols.io.6qpvr9353vmk/v1), immunohistochemistry (dx.doi.org/10.17504/protocols.io.n92ldrmd8g5b/v1), and EdU detection on sections (dx.doi.org/10.17504/protocols.io.e6nvwbdrzvmk/v1).

| **EdU reaction cocktail** |  |
| --- | --- |
| **Components** | **Volume (ul)** |
| 2 M Tris, pH 8.5 | 5 |
| 50 mM CuSO<sub>4</sub> | 2 |
| 0.5 M ascorbic acid | 20 |
| Alexa fluor azide (0.25 mg/ml) | 0.2 |
| ddH<sub>2</sub>O | 73 |
| Total | 100 |

| **Primary Antibodies** |  |  |  |  |  |
| --- | --- | --- | --- | --- | --- |
| **Antibody** | **Company, Cat#** | **RRID** | **Host species** | **Dilution** | **Antigen retrieval** |
| Anti-BrdU (Bromodeoxyuridine) | Abcam, Ab6326 | AB_305426 | Rat | 1:500 | Citric Acid |
| Anti-wide spectrum Cytokeratin | Abcam, Ab9377 | AB_307222 | Rabbit | 1:500 | None |
| Anti-ionized calcium-binding adapter molecule 1 (IBA1) | Novus Biologicals, NB100-10288s | AB_521594 | Goat | 1:400 | Tris |
| Anti-PDGFRα | R&D systems | AF1062 | Goat | 1:500 | Citrate |

| Secondary Antibodies |  |  |  |  |
| --- | --- | --- | --- | --- |
| Antibody | Company, Cat# | RRID | Host species | Dilution |
| Alexafluor 594 anti-Rabbit | Thermo Fisher Scientific, A21207 | AB_141637 | Donkey | 1:400 |
| Alexafluor 647 anti-goat | Thermo Fisher Scientific, A21447 | AB_2535864 | Donkey | 1:400 |
| Alexafluor 647 anti-rat | Thermo Fisher Scientific, A21247 | AB_141778 | Goat | 1:400 |

### Imaging and Cell Quantification

Fluorescent images were collected using Cells Sens software from Olympus on an Olympus inverted microscope (Olympus America Inc). Whole tissue sections were captured at 20X magnification and stitched automatically. Cell counting was manually performed in defined areas of the images separately in each channel using ImageJ. Percentages of positive cells were calculated by quantifying nuclei (Hoechst 33342) using the threshold tool in ImageJ.

### Statistics

Statistical analyses were performed using JMP (version Pro 15.0, SAS Institute Inc) and SPSS (version 28.0, IBM) where alpha was set at 0.05 for all statistical analyses. Graphs were made in excel and JMP. To analyze our pulse chase data across species, we performed Two-way ANOVAs for each species with EdU+, BrdU+ or double positive cells (dependent variable), and days post injury (independent variable) followed by Tukey HSD multiple comparisons to test for differences in positive cells as a function of time.

## RESULTS

### Cell cycle re-entry close to the injury site is a common response in regenerative and non-regenerative vertebrates

To assess the presence of actively cycling cells and their contribution to regeneration, we used an EdU-BrdU pulse-chase strategy (Fig. 1A-D). Uninjured animals were exposed to EdU for a 9-hr period before injury and tissue collection. Thereafter, animals were exposed to BrdU for 9-hr at either 15 or 63-hrs post injury for collection of injured tissue at 24 or 72-hrs post injury (hpi) (Fig. 1A-D). This strategy allowed us to evaluate *de novo* cell cycle entry and cell cycle progression of actively cycling cells (see methods for additional details). Next, we quantified stromal and epidermal cell proliferation in two highly regenerative amniotes with indeterminate growth (*Danio rerio* – **zebrafish** and *Ambystoma mexicanum* - **axolotl**), and in two mammals with determinate growth: the **spiny mouse** (*Acomys dimidiatus*), known for exhibiting enhanced regenerative abilities and an outbred **ND4 laboratory mouse** strain, known for having poor regenerative abilities similar to humans (Fig. 1A-D).

**Figure 1.**
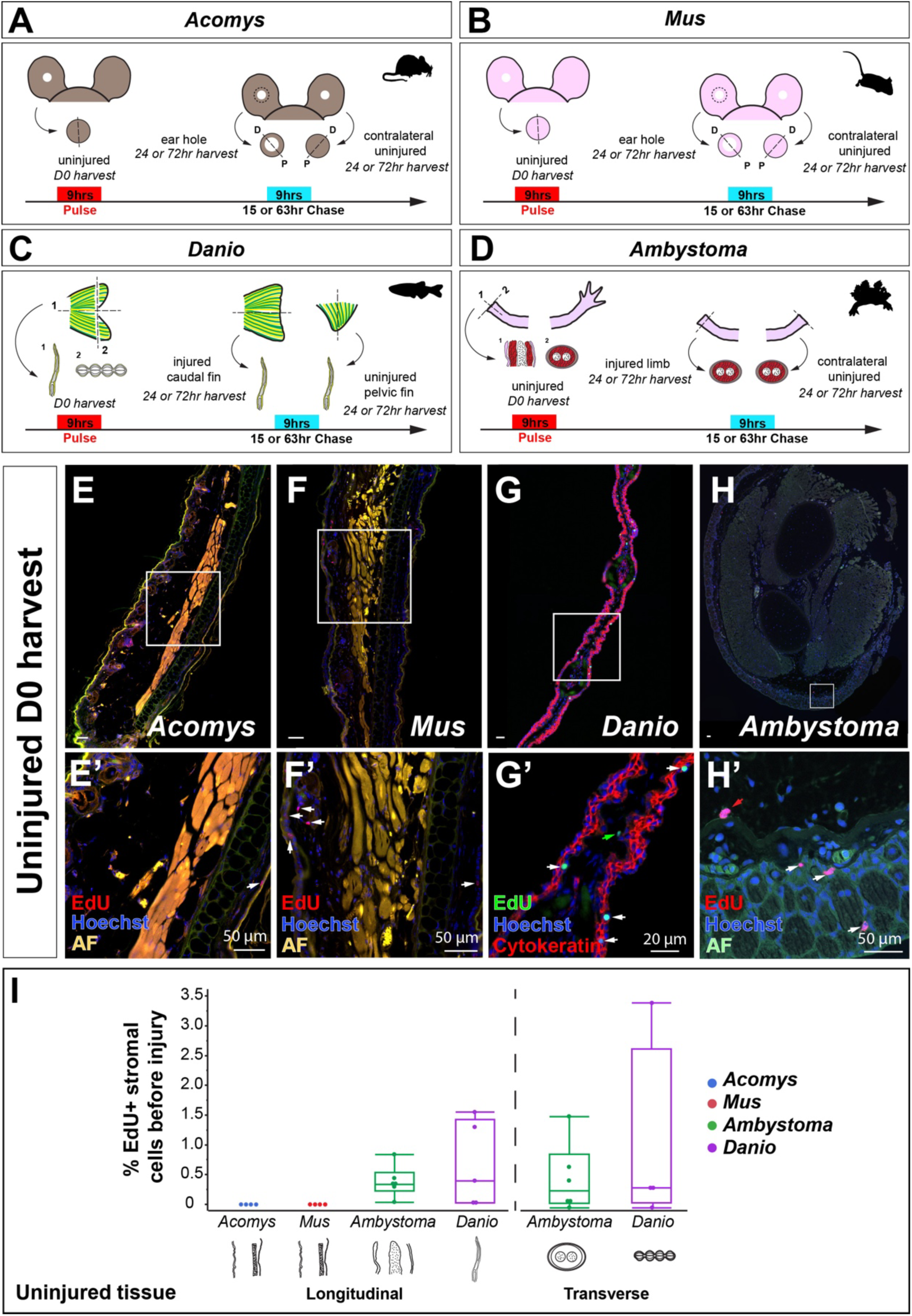
Actively cycling cells before injury in *Acomys*, *Mus*, *Danio* and *Ambystoma*. A-D) Tissue section planes and EdU/BrdU pulse-chase scheme indicating injections and harvest times; D = distal, P = proximal; 1= transverse, 2= longitudinal. E-H) Representative images of EdU+ cells in uninjured tissue (D0). Red and green arrows indicate EdU+ cells in the stromal compartment. White arrows indicate EdU+ cells in the epidermis including glands and hair follicles. I) EdU+ cell quantification as a percentage of the total number of cells in uninjured tissue from all four species. Autofluorescent (AF) tissue, including red blood cells, stains in multiple channels. Quantifications are from n = 3 biological samples in *Acomys* and *Mus*. In *Danio* (zebrafish), n = 9, In *Ambystoma* (axolotl), n = 12. See Methods for cell counting quantification. (* *p*<0.05, ** *p*<0.01).

First, we quantified EdU+ cells in uninjured connective tissue from *Acomys* and *Mus* and found only one or two cycling stromal cells across entire tissue sections prior to tissue removal (Fig. 1E’-F’, I). We did, however, find actively cycling cells among epithelial cells in the epidermis and hair follicles (Fig. 1E’-F’, Fig 4A, E). In contrast, and consistent with previous reports from *Danio* and *Ambystoma*, we found a small percentage of actively cycling cells (i.e., cells in S-phase) within uninjured connective tissue (*Danio* - 1% and *Ambystoma* - 0.5%) (Fig. 1G-I) (Johnson et al., 2018; Simkin & Seifert, 2018). These data support that cycling stromal cells are present in uninjured connective tissue from highly regenerative vertebrates with indeterminate growth modes but not in those with determinate modes of growth. Moreover, this feature does not seem to be specific to all regenerative vertebrates, as *Acomys* do not have cycling cells in uninjured stromal tissue suggesting that the absence of cycling stromal cells is not an impediment to regenerative healing.

To test whether actively cycling cells contribute to regeneration, we next quantified the number of EdU+, BrdU+ and EdU+/BrdU+ cells in stromal tissue following tissue removal. In our pulse chase scheme, EdU+ cells represent cells that were cycling prior to injury, BrdU+ cells represent those cells activated post-injury, and EdU+/BrdU+, those that were cycling prior to injury and continued cycling post-injury. At 24 hpi, *Acomys* and *Danio* showed the highest percentage of newly cycling cells (BrdU+) in response to injury (Fig. 2A, E, I, K). Although we did not observe BrdU+ cells at 24 hpi in *Mus*, newly cycling cells peaked later at 72 hpi (Fig. 2C, D, J). The percent of BrdU+ cells in *Acomys* and *Danio* also increased at 72 hpi compared to 24hrs (Fig. 2B, F, I, K). *Ambystoma* limb tissue, on the other hand, exhibited an increase in double positive cells at 72 hpi (Fig. 2G, H, L).

**Figure 2.**
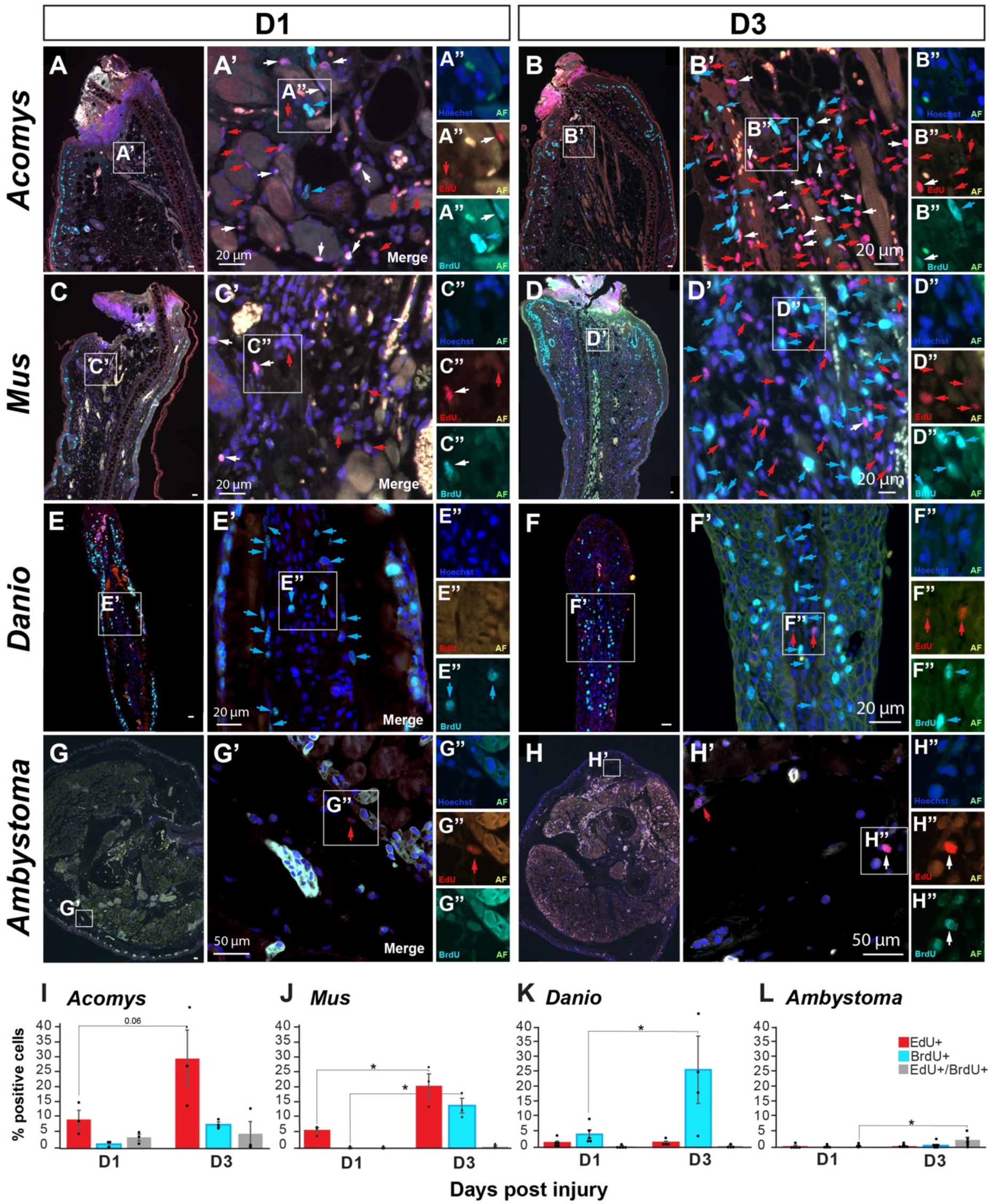
Cell cycle re-entry after injury in *Acomys*, *Mus*, *Danio* and *Ambystoma* at D1 and D3. A-H) Representative images of tissue sections after EdU/BrdU pulse-chase in *Acomys, Mus, Danio and Ambystoma* at D1 and D3. Prime images are amplifications of area of interest in whole tissue section and double prime images are amplifications of labeled cells in four channels; 405 and 488 (Hoechst and autofluorescence, AF), 594 and 488 (EdU and AF), 647 and 488 (BrdU and AF). I-L) EdU+, BrdU+ and EdU+/BrdU+ cell quantification as a percentage of the total number of cells at D1 and D3 post-injury in the four species. Quantifications are from n = 3 biological samples/ time point in *Acomys* and *Mus*. In *Danio* (zebrafish), n = 5 at D1 and n = 4 at D3, In *Ambystoma* (axolotl), n = 6 at D1, n = 6 at D3. Red arrows = EdU+ cells, light blue arrows = BrdU+ cells, and white arrows = double positive cells (EdU+/BrdU+). See Methods for cell counting quantification. (* *p*<0.05, ** *p*<0.01).

EdU+ cell counts significantly increased in *Acomys* and *Mus* at 24 and 72 hpi. However, since no EdU+ cells were found before injury, it is possible that some of these cells could be inflammatory cells (Fig. 2A-D, I, J). When EdU is injected intraperitoneally into an animal, the solution gets distributed throughout the body and incorporates into cycling cells including hematopoietic stem cells. When the animal experiences an injury, cells respond by entering the cell cycle and producing the progeny that will eventually infiltrate the injured tissue (Mescher et al., 2017). To test if the increased EdU+ cells in *Acomys* and *Mus* was due to the infiltration of inflammatory cells, we co-stained for EdU and three cell type markers (CD45, IBA1, and PDGFRα) and quantified positive cells among the EdU+ population. In *Acomys*, we found that CD45+, IBA1+ and PDGFRα+ cells represented 87%, 17.8% and 0% of the EdU positive population respectively at 24 hpi (Fig S1A-C, M) and 91.2%, 33.6%, and 4.5% respectively at 72 hpi (Fig. S1G-I, M). In *Mus* they represented 92.1%, 20.4% and 1.3% respectively at 24 hpi (Fig. S1D-F, N) and 88.7%, 44%, and 0.8% respectively at 72 hpi (Fig. S1J-L, N). Together these markers show that most EdU+ cells present in the injured site at these early time points are infiltrating leukocytes.

Together, these data show that the presence or absence of cycling stromal cells prior to injury does not predict cell cycle re-entry in the four species examined. Moreover, it documents that cell cycle re-entry is a common response to injury in regenerative and non-regenerative vertebrates with different modes of growth. Importantly, our data shows that cycling cells in uninjured tissue do not appreciably contribute to the regeneration process except for infiltrating inflammatory cells.

### Systemic cell cycle activation in response to injury is not observed in connective and epidermal tissue

To assess cell activation arising from the injury but independent of the local repair response (i.e., systemic cell activation), we collected the contralateral ear pinna in *Mus* and *Acomys*, the contralateral limb in *Ambystoma*, and the pelvic fin in *Danio* at 24 hpi and 72 hpi. Using a pulse chase strategy as described above, we looked for cell cycle re-entry in distant, uninjured tissues at 24 and 72 hpi. Although long-range cell activation in mouse satellite cells (Rodgers et al., 2014) and *Ambystoma* limb tissue (Johnson et al., 2018) has been previously reported, we did not find a significant response in the contralateral ear pinna connective tissue from *Acomys* and *Mus* (Fig. 3A-D, I, J). BrdU+ cells did not exceed 0.5% at either time point. Similarly, in *Ambystoma* and *Danio*, the number of EdU+ cells at 24 and 72 hpi was not different from the number of positive cells found in uninjured tissue at D0 and BrdU+ cells did not exceed 0.5% at either time point (Fig. 3E-H, K, L). These data indicate that a systemic response to injury is negligible in stromal tissue at 24 and 72 hpi in all four species.

**Figure 3.**
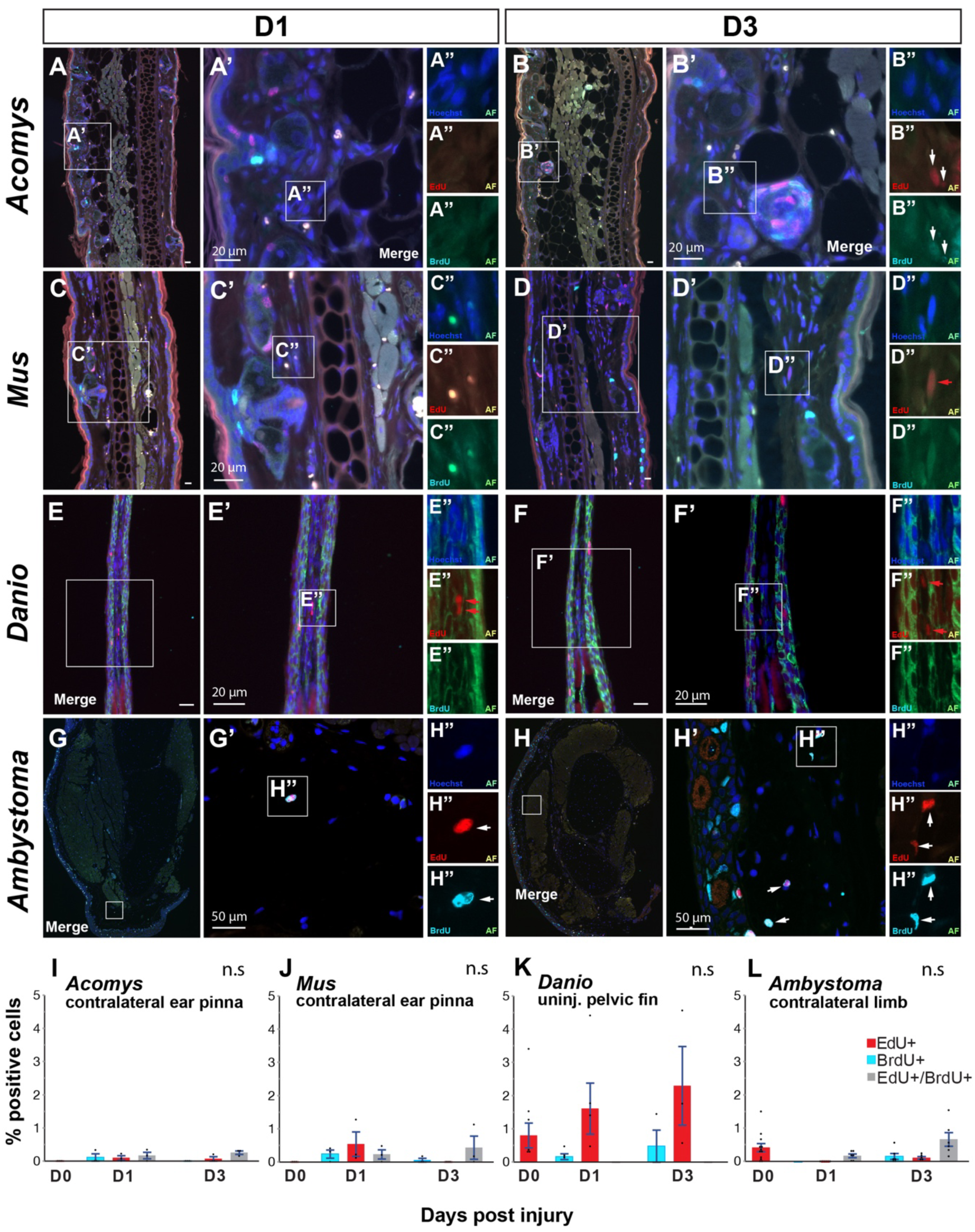
Cell cycle activation after injury in contralateral or distal uninjured stromal tissue in *Acomys*, *Mus*, *Danio* and *Ambystoma* at D1 and D3. A-H) Representative images of contralateral stromal tissue sections after EdU/BrdU pulse-chase in *Acomys, Mus and Ambystoma,* and stromal pelvic fin tissue in *Danio* at D1 and D3. Prime images are amplifications of area of interest in whole tissue section and double prime images are amplifications of labeled cells in four channels: 405 and 488 (Hoechst and autofluorescence, AF), 594 and 488 (EdU and AF), 647 and 488 (BrdU and AF). I-L) EdU+, BrdU+ and EdU+/BrdU+ cell quantification as a percentage of the total number of cells at D1 and D3 post-injury in the four species. Quantifications are from n = 3 biological samples/ time point in *Acomys* and *Mus*. In *Danio* (zebrafish), n = 5 at D1 and n = 4 at D3, In *Ambystoma* (axolotl), n = 6 at D1, n = 6 at D3. Red arrows = EdU+ cells, light blue arrows = BrdU+ cells, and white arrows = double positive cells (EdU+/BrdU+). See Methods for cell counting quantification and statistics. n.s = non-significant (* *p*<0.05, ** *p*<0.01).

Additionally, we assessed the response of basal stem cells in the epidermis to see whether this cell population responded to long-distance injury signals. We analyzed cell cycle re-entry between uninjured D0 epidermis and D1 (injured), D1 control (uninjured), and D3 control (uninjured) epidermis by comparing D0 EdU+ cells to D1 and D3 EdU+, BrdU+ and double positive cells. As expected, we found clear differences between D0 uninjured tissue and D1 injured tissue, but we failed to find differences between D0 and D1 and D3 uninjured controls in any species (Fig. 4A-T). Although not significant, we did observe a trend towards increasing BrdU+ cells in *Ambystoma* contralateral epidermis over time. Together, these data demonstrate that no systemic response to injury occurs in stromal or epidermal tissue from these species during the acute response to injury over 72hrs.

**Figure 4.**
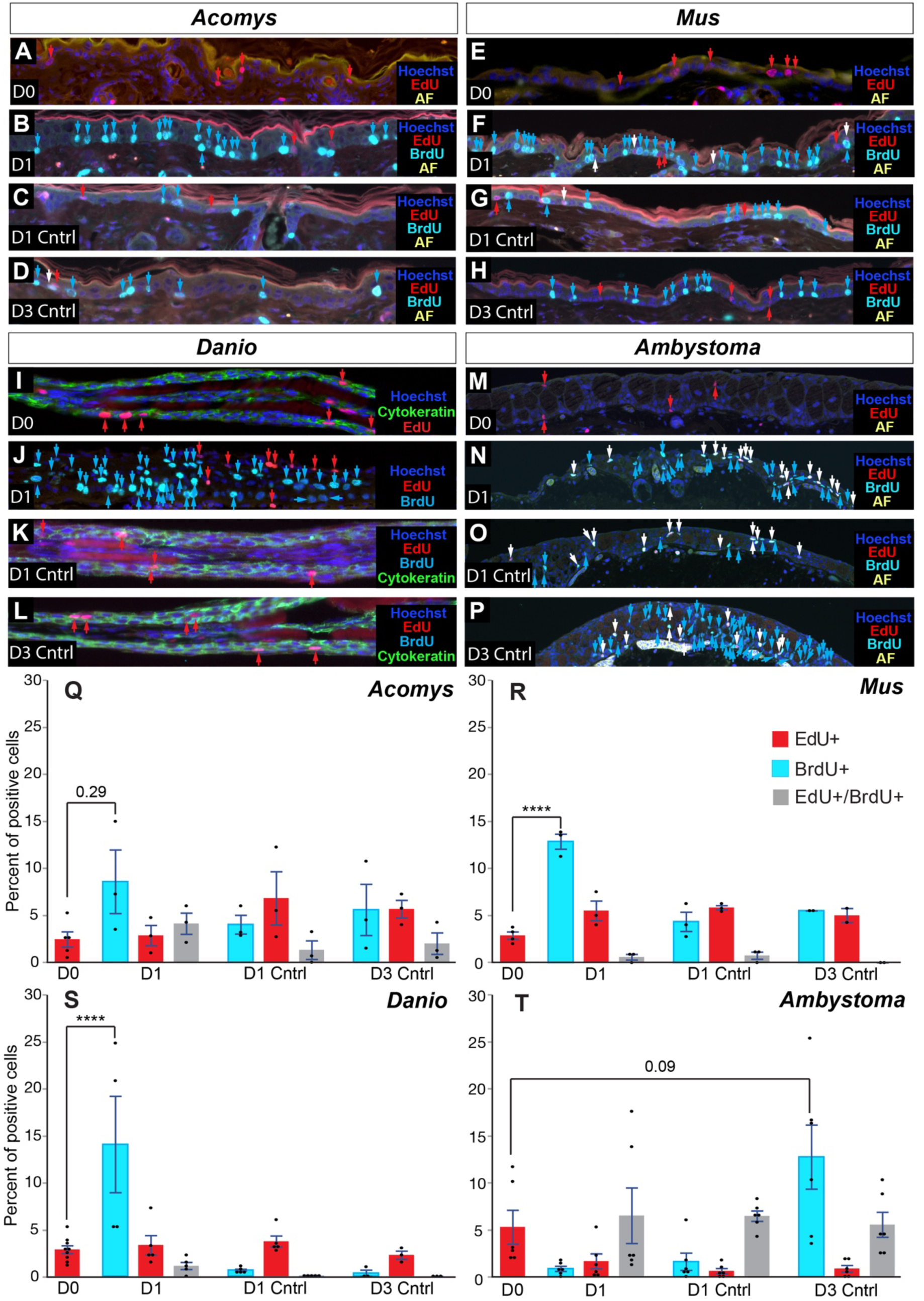
Cycling cells before and after injury in injured and uninjured contralateral or distal epidermis in *Acomys*, *Mus*, *Danio* and *Ambystoma* at D0, D1 and D3. A, E, I, M) Representative images of uninjured epidermis after a 9-hr EdU pulse in *Acomys, Mus, Danio and Ambystoma,* at D0. B, F, J, N) Representative images of injured epidermis after an EdU/BrdU pulse-chase in *Acomys, Mus, Danio and Ambystoma,* at D1. C, G, K, O) Representative images of uninjured contralateral or distal epidermis after an EdU/BrdU pulse-chase in *Acomys, Mus, Danio and Ambystoma,* at D1. D, H, L, P) Representative images of uninjured contralateral or distal epidermis after an EdU/BrdU pulse-chase in *Acomys, Mus, Danio and Ambystoma,* at D3. Q-T) EdU+, BrdU+ and EdU+/BrdU+ cell quantification as a percentage of the total number of cells at D0, D1, D1 control and D3 control in the four species. Quantifications are from n = 3 biological samples/ time point in *Acomys* and *Mus*. In *Danio* (zebrafish), n = 5 at D1 and n = 4 at D3, In *Ambystoma* (axolotl), n = 6 at D1, n = 6 at D3. Red arrows = EdU+ cells, light blue arrows = BrdU+ cells, and white arrows = double positive cells (EdU+/BrdU+). See Methods for cell counting quantification. (* *p*<0.05, ** *p*<0.01, **** *p*<0.0001).

## DISCUSSION

In this study, we observed a small population of actively cycling stromal cells present during homeostasis in indeterminate growers but did not detect a similar population in the two mammals with determinate growth. Interestingly, this pre-injury cycling population did not significantly contribute to blastema formation in *Danio*. Instead, cells that entered the cell cycle in response to injury were the major contributors to regeneration. This result supports a previous study in *Danio* where actively cycling cells marked for eight weeks before injury did not contribute to blastema formation (Nechiporuk & Keating, 2002). Importantly, our data show that cells in *Acomys* and *Mus* re-enter the cell cycle in response to injury, a result that shows cell cycle re-entry is independent of regenerative ability. Our double positive cell counts with EdU and cell type markers show that most of these cells observed early in response to injury are infiltrating leucocytes in mammals, although we did also detect a small population of cycling stromal cells. Although we did not perform such co-staining in *Ambystoma* and *Danio*, the low number of EdU+ cells in these species aligns with previous studies showing faster resolution of infiltrating cells in *Ambystoma* and *Danio* in comparison with mammals (Miskolci et al., 2019; Rodgers et al., 2020; Seifert, et al., 2012). This striking difference has been previously used as a possible explanation of the deficient regenerative abilities in mammals (Mescher et al., 2017). However, *Acomys* shows similar inflammatory cell infiltration and persistence to *Mus* suggesting other phenomena like cell phenotype or signaling are likely to explain differences in regenerative ability. While our study did not investigate cell cycle progression, an open question during complex tissue regeneration in *Acomys* is how long-term cell cycle progression and cell cycle exit is regulated so as not to create tissue overgrowth. Considered *in toto*, our cross-species analysis rejects the notion that a determinate growth mode prevents cell cycle re-entry and supports that the contribution of actively cycling cells to the blastema is minimal, at least in *Danio* and *Acomys*. A longer time course is necessary to definitively rule this out in *Ambystoma* where the cell cycle is approximately 42hrs (Wallace & Maden, 1976). Importantly, previous results comparing *Acomys* and *Mus* ear pinna tissue during regeneration and fibrotic repair respectively, shows that cell cycle re-entry in *Acomys* leads to sustained cell proliferation and regeneration whereas the upregulation of the tumor suppressors p21 and p27 prevents cell cycle progression during fibrotic repair (Gawriluk et al., 2016). Together, these data support the notion that cell cycle surveillance systems in non-regenerative species activate pathways that inhibit prolonged cell proliferation, possibly as a means to prevent tumorigenesis.

Our cross-species proliferation experiments also assessed whether cells on the other side of the animal were activated by injury (so-called systemic cell activation). This phenomenon has been reported in planarians, where an amputation causes two waves of cell cycle activation. The first one is systemic and responds to any kind of injury, while the second one is local (near the wound site) and responds to injuries where a significant amount of tissue has been lost (Wenemoser & Reddien, 2010). Similar responses have also been reported in vertebrates. Work in lab mice found that acute muscle injury elicits a systemic response in which contralaterally located stem cells (CSCs) re-enter the cell cycle and access a priming state named G_Alert_ (J. T. Rodgers et al., 2014). In contrast to quiescent stem cells (QSCs), CSCs in mouse show an increased propensity to cycle, although their numbers remain low in comparison with stem cells located at the injury site (Rodgers et al., 2014). Similarly, studies in *Ambystoma* have reported a systemic response in various tissues across the body after a limb amputation (Johnson et al., 2018; Payzin-Dogru et al., 2025).

Using contralateral control tissue collected in our experiments, we looked for primed cells *in vivo* following injury. In contrast to muscle stem cells in *Mus*, we found a negligible stromal response in *Acomys* and *Mus* with less than 0.5% of contralateral ear pinna cells responding to a systemic signal. Similarly, we could not find a significant difference between D0 and contralateral *Ambystoma* limbs or *Danio* pelvic fins at any time point. Following the same trend, we did not find a significant response in epidermal basal stem cells at either time point in any species. Contrary to our results, an *Ambystoma* study reported that contralateral limb tissue had a significant proportion of cells re-entering the cell cycle in response to long range injury (Johnson et al., 2018). However, that study quantified epidermal and stromal cells together whereas we separately quantified activated cells in stromal and epidermal tissue; a difference of approach that likely explains the different result reported across our studies. This allowed us to detect an increasing although not significant trend of cell activation in epidermal tissue by D3.

However, such activation was not evident in somatic tissue from any species studied. Together, our results show that contrary to planarians and stem cells in specific tissues or species, systemic cell cycle activation is negligible in stromal and epidermal tissue of all the vertebrates assessed here.

## ACKNOWLEDGEMENTS

We would like to thank Josh Sarli, Brennan Riddell and Ava Musarra for animal husbandry. We thank all members of the Seifert lab for helpful discussions in developing the manuscript. This research was funded in part by grants from the NIH (R01AR070313) to AWS. For the purpose of open access, the author has applied a CC BY public copyright license to all Author Accepted Manuscripts arising from this submission.

## AUTHOR CONTRIBUTIONS

EOR: Conceptualization, Data curation, Formal analysis, Investigation, Methodology, Visualization, Writing – original draft, review and editing. AWS: Conceptualization, Funding acquisition, Project administration, Supervision, Visualization, Writing – original draft, review and editing.

## AVAILABILITY STATEMENT

Data and statistical outputs are supplied in a Supplementary Data File. Protocols and key lab materials used and generated in this study are listed in the methods in a Key Resource Table alongside their persistent identifiers.

