## Supplemental figure 1 for "Actively cycling cells in uninjured connective tissue are not a prerequisite for appendage regeneration"

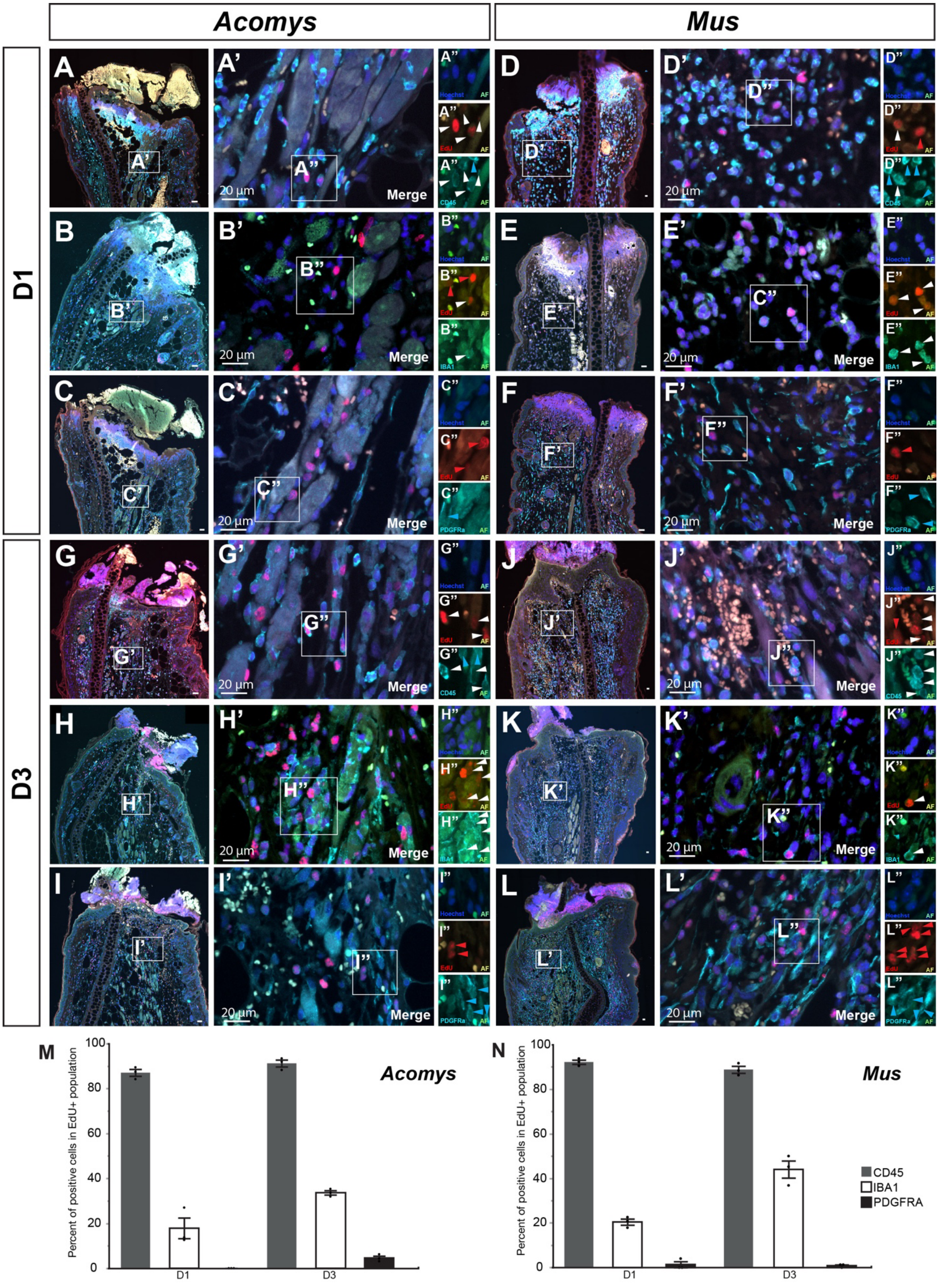
Acomys
Mus
D1
D3
Merge
20 µm
Hoechst
AF
EdU
CD45
IBA1
PDGFRa
M
N
Percent of positive cells in EdU+ population
D1
D3
CD45
IBA1
PDGFRA

**Figure S1. EdU positive cells in *Acomys* and *Mus* at D1 and D3 are mostly cycling infiltrating leucocytes.** A-L) Representative images of tissue sections double stained with EdU and CD45, IBA1 or PDGFRα in *Acomys* and *Mus* at D1 and D3. Prime images are amplifications of area of interest in whole tissue section and double prime images are amplifications of labeled cells in four channels; 405 and 488 (Hoechst and autofluorescence, AF), 594 and 488 (EdU and AF), 647 and 488 (Cell type marker and AF). M-N) CD45+, IBA1+ and PDGFRα+ cell quantification as a percentage of the total number of cycling cells (EdU+) at D1 and D3 post-injury. Quantifications are from n=3 biological samples/ time point in *Acomys* and *Mus*. Red arrows = EdU+ cells, light blue arrows = BrdU+ cells, and white arrows = double positive cells (EdU+/BrdU+). See Methods for cell counting quantification. (* $p<0.05$, ** $p<0.01$).
